# Age- and sex-dependent sibling effects on early-life survival in preindustrial humans

**DOI:** 10.1101/2025.06.30.661956

**Authors:** Mark Spa, Euan A Young, Virpi Lummaa, Erik Postma, Hannah L Dugdale

## Abstract

Siblings are an important part of an individual’s early-life environment and may therefore play an important role in shaping an individual’s fitness. The quantification of sibling effects is challenging, especially in long-lived species with extended parental care and overlapping generations, such as humans. Here we quantify how the survival status, age, and sex of older siblings shape childhood survival across 2941 focal individuals born between 1750–1870 using historical parish data from Switzerland. While the total number of older siblings was not associated with an individual’s childhood survival, distinguishing between siblings by their survival status, age, and sex revealed several associations, which in some cases also interacted with the sex of the focal individual: While older brothers close in age reduced the survival of girls but not boys, having more older sisters born close in age improved their younger sibling’s survival. Our results, therefore, show that sibling interactions play an important role in shaping early-life survival, and highlight that the strength and direction of these effects are context-dependent and can arise through biological and cultural factors. We encourage future studies on sibling interaction to consider siblings’ survival, age and sex, in both humans and other species.

## INTRODUCTION

How the early-life environment shapes individual variation is of keen interest to evolutionary ecologists (Grafen, 1988; Lindström, 1999). Across a variety of animal species, older siblings are one important aspect of the early-life environment (Emlen, 1995; Hudson & Trillmich, 2008) with the potential to shape evolutionarily and demographically important traits, such as survival and reproduction (Clutton-Brock, 1988). Thus, determining whether sibling interactions are cooperative or competitive is important for understanding whether siblings promote or constrain species’ optimal brood, litter, or family sizes. However, quantifying the effects of sibling interactions on fitness components has proved challenging, especially in species with long, slow life-histories and overlapping generations (Hudson & Trillmich, 2008).

Whether siblings provide benefits or compete depends upon a species’ life-history and degree of cooperation. When siblings do not act cooperatively, having more siblings will lead to competition because a limited amount of resources, such as parental care, is split among more offspring (Lack, 1947; Trivers, 1974). However, competitive abilities amongst siblings are not always equal, leading to variation in how siblings interact. In species where offspring are not produced simultaneously, older siblings will be more developed and have competitive advantages over younger siblings (Trillmich & Wolf, 2008). Competitive abilities between siblings can also vary by sex in species that are sexually dimorphic (Clutton-brock, 2016). Competitive sibling interactions may then be minimised or reversed by cooperative behaviours, promoted through kin selection (Hamilton, 1964; Parker et al., 1989), such as direct help or behaviours that increase shared familial resources (Hertwig et al., 2002). Thus far, ecological studies have largely focused on within-brood sibling interactions in birds (Hudson & Trillmich, 2008) and understanding the balance of cooperative and competitive sibling interactions across sexually dimorphic species with multiple reproductive events is a place of active research [see (Berger et al., 2021) for elephants and (Fox et al., 2017) for humans].

In humans, the impact of sibling interactions has attracted interest from researchers from different fields, studying a variety of cultures and traits, including dispersal behaviour (Beise, 2004; Zhou et al., 2022; Nitsch et al., 2013), nutritional status (Hagen & Barrett, 2009; Helfrecht & Meehan, 2016), educational attainment (Blake, 1989; Steelman et al., 2002), and marital timing (Suanet & Bras, 2014). Some studies have also looked at fitness outcomes in preindustrial societies, namely childhood survival (Borgerhoff Mulder, 1998; Fox et al., 2017; Nitsch et al., 2013; Marco-Gracia & López-Antón, 2025). In these human societies, short interbirth intervals combined with long developmental times (Harvey & Clutton-Brock, 1985; Kaplan, 1997) result in parents raising multiple dependent children simultaneously (Kramer, 2005; Sear & Mace, 2008). While humans are a highly cooperative species, older siblings that are closer in age and still dependent on parents could be expected to compete with younger siblings and negatively impact their development (Kramer, 2002). This might occur through direct competition for parental resources, or via effects on the health and condition of the mother during previous pregnancies (Boerma & Bicego, 1992; Whitworth & Stephenson, 2002). Conversely, it is expected that older and more independent siblings (often defined as being at least 5 years older; e.g., Riswick & Hsieh, 2020) can have positive effects on survival through cooperative behaviours (Kramer, 2002). These behaviours can involve direct childcare (Kramer, 2005), or be indirect: older siblings may take over tasks that allow parents to focus on caring for newborns (Riswick & Hsieh, 2020), or increase familial resources [e.g. through foraging, agricultural labour (Kramer, 2005; Kramer & Boone, 2002), or paid labour in industrialised societies (Nag et al., 1978)]. Although isolating specific mechanisms is challenging, studies have generally found findings in support of these predictions: more older siblings close in age are associated with lower early-life survival (Riswick & Hsieh, 2020), while more older siblings further in age are associated with higher early-life survival (Crognier et al., 2001; Crognier et al., 2002).

Biologically and culturally mediated sex differences (Frayer & Wolpoff, 1985) can interact with sex differences in development time to make sibling interactions in humans highly sex-dependent (Rickard et al., 2009). Biologically, because males are on average larger and require more resources (Thurstans et al., 2022, Invernizzi et al., 2024), the effect of having older brothers on their younger siblings’ survival could be expected to be more negative compared to the effect of older sisters (Rickard, 2008). However, males also have lower childhood survival rates than females, owing perhaps to their weaker immune function (Thurstans et al., 2022; Bawkin, 1929; Drevenstedt et al., 2008). The death of a previously born older sibling may decrease mortality risks of the younger sibling (Boerma & Bicego, 1992), indicating that the survival status of older brothers can moderate their effects, and that, in addition to age differences, both the sex and survival status of siblings is important to account for in studies.

The negative biological effects of having older brothers can in turn be moderated by cultural factors, which could explain why several studies found no sex-specific effects of older siblings on early-life survival (Crognier et al., 2001; Crognier et al., 2002; Sear, 2008). For example, having more older brothers enhanced survival to 15 years in preindustrial Finland, possibly through providing economic contributions (Nitsch et al., 2013). On the other hand, in cultures where only sisters provide help to younger siblings, the number of older sisters positively correlates with survival (Sear et al., 2002). Finally, in cultures with a preference for sons (e.g. due to patriarchal inheritance), sisters may be more negatively affected by the presence of brothers (Riswick & Hsieh, 2020). This may happen because more resources are allocated to males rather than direct harm by parents or siblings (Marco-Gracia & Beltrán Tapia, 2021; Rosenblum, 2015), but, regardless, the effect can be so strong that it reverses the biological differences in childhood mortality between sexes (Fuse & Crenshaw, 2006). Son preference can, in turn, increase the competition for resources among sisters, placing younger female siblings at an even greater disadvantage (Muhuri & Menken, 1997). When this combines with older brothers bringing in more resources, it can feed into a wider picture of same-sex competition but opposite-sex benefits (Fox et al., 2017; Riswick & Hsieh, 2020; Voland & Dunbar, 1995). It is thus crucial that studies account for age differences, survival statuses and how they interact with sex when examining the effects of siblings on early-life survival.

Few studies have simultaneously considered both siblings who may compete for resources and those who may provide support (but see Riswick & Hsieh, 2020). Here, we aim to fill this knowledge gap and examine evidence for cooperative and competitive interactions between siblings shaping an important factor in the evolutionary and demographic history of humans: childhood survival (Volk & Atkinson, 2013), defined here as survival to age 5. To this end, we use historical life-history data from 2941 individuals, born 1750-1870, adapted from Swiss church parish records (Kubly-Müller, 1912). In this population, during this period, both fertility and childhood mortality were high, making the population particularly valuable for investigating the effect of siblings on childhood survival patterns. We first quantify the association between childhood survival and total number of older siblings controlling for potentially important confounders, such as grandparental presence (Chapman et al., 2021), parental presence (Atrash, 2011), parental age (Arslan et al., 2017), and socioeconomic status (Braudt et al., 2019). We then conduct a decomposition of the number of older siblings in a series of models, separating siblings by whether siblings had died before the birth of the focal individual (their survival status), whether they were born close (< 5 years) or far in age (≥5 years) from the focal individuals, and whether they were sisters or brothers. We estimate the associations between childhood survival and the number of older siblings in each of these categories, while allowing for these associations to be dependent on the sex of the focal individual. On the whole, these analyses provide a uniquely detailed insight into the consequences of sibling interactions, and how these are modulated by cultural factors such as parental preferences or gender-specific roles within the family.

We predict that the number of older siblings that are ≥5 years of age increases survival because the positive effects of helping outweigh the negative effects of competition. Conversely, we expect the number of older siblings that are <5 years of age to have detrimental effects, owing to parental health and/or competition for parental resources. We also expect older sibling effects to vary based on the sex of both the older siblings and the focal individual and predict brothers to have more detrimental effects owing to larger resource requirements. Finally, a son preference would manifest itself as an interaction between the sex of the sibling and the sex of the focal individual, with the effect of the number of brothers being more negative (or less positive) for females than males.

## METHODS

### Study population

We used data from an extensive genealogical archive (Kubly-Müller, 1912) which covers two parishes situated on the Swiss Alpine plateau: Linthal (46°55’ N 9°00’E) and Elm (46°55’ N 9°10’E). This archive contains birth (including unbaptised individuals), marriage, and death dates for individuals born between 1540 and 1998, but 73% were born after 1800. For 96% of the individuals, both their birth date and the identity of their parents were known, allowing for the characterisation of family structure at birth. Although the death date was missing for 41% of individuals, this was mainly due to emigration as an adult and deaths before the age of 5 were unlikely to have been missed.

We limit our analyses to individuals born between 1750 and 1870 as sample sizes for earlier years were relatively small (e.g., less than 30 recorded births per year). After 1870, early-life survival gradually improved in Switzerland (Siegenthaler & Ritzmann-Blickenstorfer, 1996). During this period, the median lifespan was 31, and survival from birth to age 5 (childhood survival) was relatively low (67%). These values are broadly consistent with historical estimates from other 18th and 19th European populations (Roser, 2023; Volk & Atkinson, 2013). At the same time, fertility was high, with a median of five children per reproductive woman, ranging from 1 to 22. Hence, this can be considered a stage one demographic transition (i.e. pre-industrialised) population (Chesnais, 1992; Corbett et al., 2018). The population is furthermore representative of a northern or western European population during this period, with relatively late ages-at-first birth (median age 25) owing to the wealth accumulation that was required pre-marriage (Lee, 2003). During this period, the region can be considered pre-industrial, as by 1850 around 50% of the Swiss population was agricultural (Rachoud-Schneider, et al., 2007), and in our data 41% of children had a father who worked in the agricultural sector (n = 2849/6987).

### Childhood survival

We treat childhood survival as a binary variable defined as survival until age 5 (Riswick & Hsieh, 2020, Lawson et al., 2012; World Health Organisation, 2022). Survival was determined for all individuals that had a recorded birth and death year. This included 194 individuals who died on the day they were born. Individuals with an unknown year of death that were known to have married and/or reproduced were assumed to have survived beyond the age of 5 (n = 2328). Overall, childhood survival status could be determined for 93% of the individuals with a known birth date (n =11878).

### Sibling classifications

Using the identity of their parents, we grouped individuals into families and counted the number of older siblings at an individual’s birth, focusing solely on full siblings of parents that only married once, which ranged from 0 to 15 older siblings. In addition to the total number of older siblings, for all older siblings with known birth and death dates, we determined their survival status until the birth of the focal individual (living older siblings range = 0 - 11, deceased older siblings range = 0 - 8). We also determined if these older siblings were less than (range = 0 - 4) or at least five years (range = 0 - 10) older than the focal individual (hence <5 or ≥5, respectively). These categorisations aimed to distinguish between siblings that, for the majority of the focal individual’s first five years, were unlikely (<5 years older) or likely (≥5 years older) to have been able to provide benefits (see Crognier et al., 2001; Riswick & Hsieh, 2020). These may include direct help or care, but also a contribution to family wealth through labour (Minge-Kalman, 1978). Finally, we distinguished between male and female older siblings in each age category (i.e., brothers and sisters) (brothers <5 older: range = 0 - 4; brothers ≥5 older: range = 0 - 7; sisters <5 older: range = 0 - 3; sisters ≥5 older: range = 0 - 6).

### Focal individual characteristics

For each focal individual, their sex was recorded (males: n = 6567; n = 6243 females) to account for factors such as the higher susceptibility of males to mortality during early life and son preference (Bawkin, 1929; Drevenstedt et al., 2008; 2012; Marco-Gracia & Beltrán Tapia, 2021). This also allowed for the examination of sex-dependent associations between childhood survival and the number of older siblings (see *Statistical analyses*), as found in other studies (Nitsch et al., 2013). Accounting for all older siblings also automatically controls for potential effects of being firstborn on survival and we therefore did not separately incorporate a firstborn variable (Faurie et al., 2009). We excluded twins from the analyses as focal individuals (n=257) owing to the differences in survival and other factors associated with twins (Lummaa, 2001; Rickard et al., 2022).

### Parental variables

Following Evans et al. (2018), we used the father’s occupation as a proxy for a family’s socioeconomic status. Occupations were standardised following the Historical International Standard Classification of Occupations (HISCO) (Leeuwen et al., 2002) and assigned a numeric value for socioeconomic status using the historical social stratification scale (HISCAM, Lambert et al., 2013). HISCAM uses records of intergenerational interactions and marriages between different occupations from 1800–1938 across northern Europe and Canada to assign different occupations a socioeconomic status ranging from 1-100 (Lambert et al., 2013). In cases where multiple occupations were present (n = 472 and as many as 8), we used the occupation with the highest HISCAM. A HISCAM score was assigned to 2333 fathers with recorded occupations, providing measures of socioeconomic statuses that ranged from 39.9 to 99 on an interval scale (servant to lieutenant, respectively), resulting in 7438 focal offspring (58%) with a known family socioeconomic status that would be used to control for potential positive effects of wealth and social status on childhood survival (Pettay et al., 2007).

We determined whether the mother and/or father died during the focal individual’s first 5 years (918 and 524 cases, respectively) to control for the negative impact this may have on offspring survival (Li et al., 2022, Moucheraud et al., 2015). Additionally, both the mother’s and father’s age at birth were included to account for potential parental age effects on early-life survival (Gillespie et al., 2013; Arslan et al., 2017). Finally, we included the number of grandparents alive in the first 5 years of the child to control for the positive effect this may have on their grandchildren’s survival (Chapman et al., 2021).

### Statistical analyses

We modelled the association between the number of older siblings and childhood survival using generalised linear mixed models (GLMMs) with the *glmer* function from *lme4 1.1.31* (Bates et al., 2015) in *R 4.2.2* (R core team, 2022), with a binomial error and logit-link. We first ran a baseline model (*m1*) estimating the association between childhood survival and the number of older siblings in total. We then ran separate models increasing in complexity, decomposing the number of older siblings into further categories. First, we split the number of older siblings into those that were alive or deceased at the focal individual’s time of birth (*m2*). We then decomposed these categories further into the number of living or deceased older siblings born close (<5 years) and far in age (≥5 years) from the focal individual (*m3*). Finally, we further decomposed these categories into the number of older living and deceased brothers and sisters born close (<5) and far (≥5) in age (*m4*).

All models were conditioned upon the following categorical fixed effects: the sex of the focal individual, the birth parish (either Linthal or Elm), and if the focal individual experienced the death of their mother or father during childhood. As linear covariates we included mother and father age at the focal individual’s birth, socioeconomic status, and the number of grandparents alive at the date of birth of the focal individual. Squared terms were also added for parental age effects, but removed, least significant first, if non-significant to aid interpretation of the first-order effects. We modelled variation in childhood survival among families and across 5-year parish-specific birth cohorts by including both as random effects. Additionally, we fitted random slopes for each sibling variable to quantify their effect within families (Schielzeth & Forstmeier, 2009). We also tested for interactions between the sex of the focal individual and all variables relating to the numbers of siblings of different categories in all models to test if any sibling interactions were sex dependent. Models were fitted using only individuals informative for all predictors (N=2941; 1454 females and 1487 males).

Significance of all fixed effects was determined using likelihood ratio tests (LRTs) with the Chi-squared distribution, using the *drop1* function (*stats 4.2.2*, R core team, 2022). Interactions were removed if non-significant (stepwise, highest p-values first) to improve interpretability of the results. Results including non-significant interactions for each model are shown in tables S2, S5, and S6. If an interaction with the focal individual’s sex was significant, a post hoc test was conducted using *emmeans 1.8.4-1* (Lenth, 2023) to determine if the association between childhood mortality and the variable was significant within each sex. DHARMa *0.4.6* was used for model diagnostics (Hartig, 2020), specifically, the KS test and QQ plots were used to examine whether residuals followed a normal distribution, and tests of overdispersion, homoscedasticity, and for outliers were performed. These tests revealed no violations. Collinearity between variables was low for most variables across all models (VIF < 5) and only surpassed 4 for the maternal age variable (assessed using *vif* from *car 3.1.1,* Fox et al., 2019)). Other packages used were: *ggeffects 1.1.5,* to predict the differences in survival between children with different numbers of siblings presented in the main text with all other predictors held at their reference for categorical predictors or the mean for numeric predictors (Lüdecke, 2018), and for data visualisation, *ggplot2 3.4.1* and *ggpubr 0.5.0* (Wickham, 2016; Kassambara, 2020). To aid model convergence, the "bobyqa" optimizer was used, and all non-categorical predictor variables were mean-centered and scaled to a standard deviation of 1.

## RESULTS

In total, 73% of all individuals in our study survived childhood (n = 2141/2941). In our baseline model (*m1*), an individual’s childhood survival was not associated with how many older siblings they had (0.91 [0.77 - 1.08], p = 0.282, Figure 1, Table S1). This association did not vary across families (p = 0.157, Table S1) and was not dependent on the sex of the focal individual (p = 0.551, Table S2). However, childhood survival was higher for individuals born in Elm than in Linthal (0.674 [0.527 - 0.861], p = 0.003, Table S1), for individuals with mothers of intermediate age and mothers who survived the first five years of their life (0.891 [0.823 - 0.965], p = 0.005, and 1.923 [1.068 - 3.461], p = 0.032, respectively, Table S1), and for individuals with older fathers (1.206 [1.022 - 1.422], p = 0.027, Table S1). Childhood survival did not associate with paternal survival across the first five years, the families’ socioeconomic status, the sex of the focal individual, or the number of grandparents alive at birth (p > 0.05, Table S1) but showed significant variation across families but not birth cohorts (p < 0.001 and p = 0.120, respectively, Table S1).

**Figure 1.**
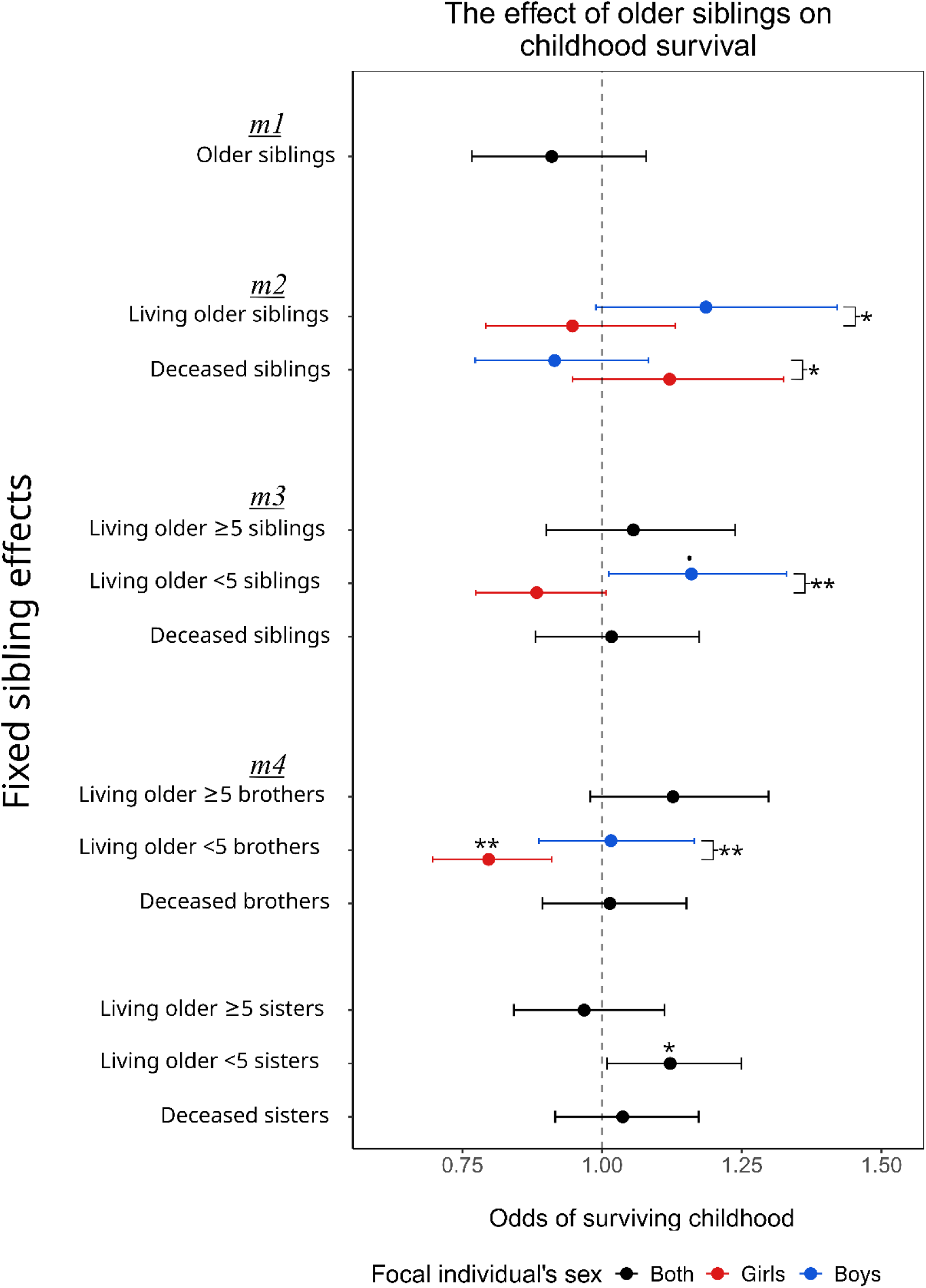
Associations between the total number of older siblings and childhood survival (*m1*) and then when decomposed by their survival status (*m2*), age (*m3*), and by sex (*m4*), in a preindustrial Swiss dataset. On the x-axis are the respective odds ratios of surviving and error bars show the 95% confidence intervals. P-values from LRT tests are shown as: • = p < 0.1, * = p < 0.05, and ** = p < 0.01. We present sex-specific odds ratios and confidence intervals (females = red; males = blue) when associations were sex-dependent.

However, dividing focal individuals by their sex and their older siblings according to whether they were alive when the focal individual was born (*m2*), revealed associations between childhood survival and the total number of both living and deceased older siblings that were dependent upon the sex of the focal individual (p = 0.018 and p = 0.026, respectively, Table S3, Figure 1). For example, having two living older siblings versus zero increased survival from 73% [51% - 88%] to 76% [56% - 89%] for males, but decreased the survival probability of females from 79% [60% - 90%] to 78% [0.58% - 0.90%]. Conversely, having two deceased older siblings versus none was associated with a decrease in survival in boys, from 78% [57% - 90%] to 75% [53% - 88%], but an increase in girls from 77% [56% - 90%] to 80% [61% - 91%]. However, within boys and girls, the effect of the of (living or deceased) older siblings was not statistically significant (the number of living older siblings in boys, p = 0.127, and girls, p = 0.794; the number of deceased older siblings in boys, p = 0.515, and girls, p = 0.334, Figure 1).

We then further divided the number of living older siblings into those close in age (<5 years old when the focal individual was born) or far in age (≥5 years of age) (*m3*, Table S4). In this model, the number of deceased older siblings and living older (≥5) siblings did not affect childhood survival (1.017 [0.881 - 1.174], p = 0.817, and 1.056 [0.900 - 1.238], p = 0.516, respectively, Table S4, Figure 1) and these associations were not dependent upon the sex of the focal individual (p = 0.080 and p = 0.386, respectively, Table S5). The association between childhood survival and the number of living older (<5) siblings was however dependent on the sex of the focal individual (p = 0.003, Table S4): Having two living older (<5) siblings versus none, increased the survival of boys from 73% [51% - 88%] to 80% [60% - 91%], but decreased the survival of girls from 80% [61% - 91%] to 75% [53% - 89%]. However, overall, the survival of boys and girls separately was not significantly affected by the number of living older (<5) siblings (in boys, p = 0.065, and girls, p = 0.124, Figure 1).

The cause of the sex-dependent association between childhood survival and the number of living older (<5) siblings was revealed when decomposing the number of living older siblings close and far in age into brothers and sisters (*m4*). Here, we found a sex-dependent association between childhood survival and the number of older (<5) brothers (p = 0.007, Table 1): Girls having, for instance, two rather than no older (<5) brothers had a survival probability of 69% [44% - 86%] versus 82% [63% - 0.92%] (p = 0.002, Figure 1), but boys’ survival was not associated with the number of older (<5) brothers (77% [55% - 90%] versus 77% [55% - 91%], p = 0.966, Figure 1). Individuals with more living older (≥5) brothers had marginally higher survival (p = 0.105, Table 1, Figure 1) such that the survival probability of individuals with two living older (≥5) brothers versus none increased from 78% [58% - 91%] to 82% [63% - 93%], regardless of their sex (p = 0.140, table S6). Having more living older (<5) sisters was also positively associated with an individual’s childhood survival (p = 0.029, Table 1, Figure 1) such that the survival probability of individuals with two older (<5) sisters versus none increased from 78% [58% - 91%] to 84% [66% - 94%], regardless of the focal individual’s sex (p = 0.124, Table S6). However, childhood survival was not associated with the number of living older (≥5) sisters, or the number of deceased older brothers or sisters (p = 0.656, p = 0.831, and p = 0.562, respectively, Table 1, Figure 1). The associations of the sex- and age-specific sibling variables with childhood survival showed no variation across cohorts (p > 0.05, Table 1), but there remained unexplained variation in childhood survival between families (p < 0.001, Table 1)

**Table 1.**
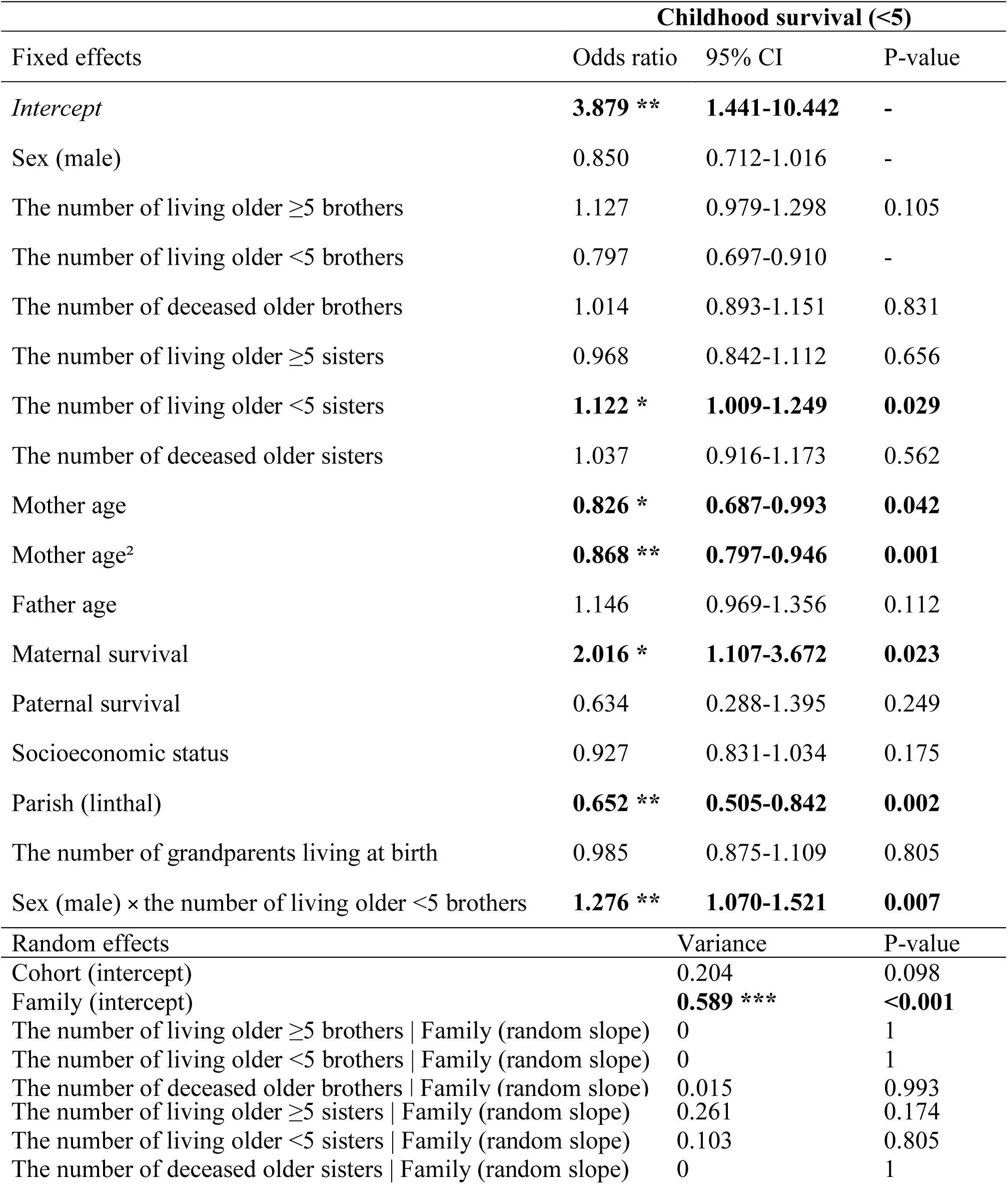
Logistic GLMM showing the association between childhood survival and the number of older siblings from *m4* (N = 2941), in a preindustrial Swiss dataset. Here, we have decomposed the number of older siblings into deceased and living older sisters and brothers born close and far in age (≥5) the focal individual and present their associations with childhood survival upon the other factors. Odds ratios, 95% confidence intervals, variation explained by random effects and p-values from LRT tests are presented. Interactions are shown with a multiplication symbol (×).

The associations between childhood survival and all other variables largely remained consistent across models (*m1*-*m4*, Table 1 and S1-S6) with only the association between father age and childhood survival being non-significant in *m2* (p = 0.088; Table S3), m3 (p = 0.098; Table S4) and m4 (p = 0.112; Table 1).

## DISCUSSION

Although at first sight childhood survival did not appear to be associated with the total number of older siblings (Figure 1), distinguishing between siblings on the basis of their survival status, sex, and age difference revealed both positive and negative associations. Thus, these results argue against exclusively positive effects of older siblings on survival (Marco-Gracia & López-Antón, 2025). Instead, they suggest that some siblings provide benefits and others are detrimental, and that these also depend on the sex of the focal individual. Thereby we show that siblings interactions are an important – but complex and context-dependent – component of the early-life environment.

Our analyses revealed several results consistent with a parental preference for sons. First, we found that girls – but not boys – with more living older brothers close in age had reduced childhood survival. Having older brothers that are close in age could be detrimental to younger siblings of either sex due to their larger size and greater energetic demands (Thurstans et al., 2022; Invernizzi et al. 2024; Rickard, 2008), but as this cost was limited to females, it suggests that parents might have tried to shield their younger sons from these costs, consistent with the male-favoured or gendered resource dilution model (Kalmijn and van de Werfhorst 2016; Riswick & Hsieh, 2020). We also found that having more living older siblings increased childhood survival of boys more than of girls, whereas having more dead older siblings benefitted females more. Although these sex-dependent interactions became weaker with further decomposition (Tables S4, S6, and 1), this suggests that any help provided by siblings benefitted boys more than girls, and that deaths caused net benefits for sisters. Evidence of male preference is in line with the primogeniture patriarchal inheritance in our study system (Hobson, 1929). However, unlike other studies (Fox et al., 2017; Nitsch et al., 2013; Riswick & Hsieh, 2020), we did not find lower childhood survival of those born with many same-sex older sibling, and hence no evidence for same-sex sibling competition within a human family (Table 4).

Contrary to the number of older brothers close in age, the number of older sisters close in age was positively associated with the survival of their younger brothers and sisters. This differs from previous studies, which have found either positive associations between the total number of older sisters for only males but negative for females – perhaps due to same-sex competition (Riswick & Hsieh, 2020) – or a positive effect of the total number of older sisters on male survival but no effect on female survival (Nitsch et al., 2013). Other studies (Nitsch et al., 2013; Riswick & Hsieh, 2020) have suggested this could be a signal of older sisters helping younger siblings, but this cannot explain why we did not find similar associations for older sisters more distant in age. Alternatively, individuals with more older sisters (<5) would less likely be exposed to older brothers (<5) and do not suffer the negative consequences of being a firstborn (Björkegren and Svaleryd, 2023), which would also give rise to positive associations between the number of older sisters close in age and survival. However, firstborn effects on survival are nuanced (Faurie et al., 2009) and, overall, these associations illustrate that while siblings have detectable effects on early-life survival, these are difficult to interpret and likely multifaceted.

Albeit weak, we also found some evidence that older brothers further away in age helped younger siblings survive (Figure 1). No such effects were found for older sisters. This is similar to associations found by Nitsch et al. (2013), who hypothesised that older brothers might have helped the productivity of family farms through providing labour. Future studies could test this by comparing farm-owning vs non-farm-owning families. In our population, there was also a strong wage gap between males and females meaning older brothers may have benefited their families’ resources through income, while older sisters had a more limited ability to do so (Davatz, 1980). Further, as potential inheritors, older brothers may also have stayed with their families for longer than sisters, giving them greater opportunity to influence the development of their younger siblings than older sisters. Finally, the absence of any associations between the number of dead brothers and childhood survival further argues for a causal effect of living brothers’ behaviour benefitting younger siblings, although the precise mechanism remains unknown.

Our study has some limitations. Firstly, its observational nature makes causal inference challenging, especially when the underlying mechanisms of sibling interactions are not well understood. Isolating sibling interactions in GLMMs (or comparable approaches such as cox-proportional hazard models) in human populations is often challenging and other variables (e.g., parental age) may capture birth order effects. To better account for these complexities, future studies could explore using structural equation models or event history analysis (Singer & Willett, 2003). Secondly, although we found evidence of older brothers and sisters influencing the early-life survival of their younger siblings, these effects were close to the threshold of statistical significance. This illustrates the difficulties of disentangling these effects in historical human populations, even with very large sample sizes.

In this study, we provided a comprehensive decomposition of how older siblings shape survival in early life. We emphasise the need to consider interactions with the sex, survival, and age of siblings, which are mediators of both the strength and the direction of sibling effects. Overall, we provide a rare insight into how sibling interactions shape survival in a long-lived species, revealing signals consistent with both cooperative and competitive interactions mediated by biological and cultural factors. Thereby these results show that siblings are an important component of the early life environment.

## Supporting information

All supplementary tables

## ACKNOWLEDGEMENTS

We thank Beat Mahler and Fritz Rigendinger of the Landesarchiv des Kantons Glarus for enabling access to the data. We thank Aïda Nitsch and the Dugdale Research Group for feedback throughout the project and Maaike A. Versteegh for comments on the writing. but

## DATA AVAILABILITY STATEMENT

Code and data necessary for reproducing results are available privately for reviewers at: https://dataverse.nl/privateurl.xhtml?token=f65e6334-203d-4574-8b7b-ff336eeff5ff

## REFERENCES

Arslan, R.C., Willfuhr, K.P., Frans, E.M., Verweij, K.J.H., Burkner, P.-C., Myrskyla, M., Voland, E., Almqvist, C., Zietsch, B.P. & Penke, L. (2017) Older fathers’ children have lower evolutionary fitness across four centuries and in four populations. Proceedings of the Royal Society B: Biological Sciences. 284 (20171562), 1–9. doi:10.1098/rspb.2017.1562.

Atrash, H. K. (2011). Parents’ death and its implications for child survival. Revista brasileira de crescimento e desenvolvimento humano, 21(3), 759.

Bates, D., Mächler, M., Bolker, B.M. & Walker, S.C. (2015) Fitting linear mixed-effects models using lme4. Journal of Statistical Software. 67 (1). doi:10.18637/jss.v067.i01.

Bakwin, H. (1929). The sex factor in infant mortality. Human Biology, 1(1), 90.

Beise, J. (2004) The helping and the helpful grandmother - The role of maternal and paternal grandmothers in child mortality in the 17th and 18th century population of French Settlers in Quebec, Canada.p.WP-2004–004. doi:10.4054/MPIDR-WP-2004-004.

Berger, V., Reichert, S., Lahdenperä, M., Jackson, J., Htut, W. & Lummaa, V. (2021) The elephant in the family: Costs and benefits of elder siblings on younger offspring life-history trajectory in a matrilineal mammal. Journal of Animal Ecology. 90 (11), 2663–2677. doi:10.1111/1365-2656.13573.

Björkegren, E., & Svaleryd, H. (2023). Birth order and health disparities throughout the life course. Social Science & Medicine, 318, 115605. 10.1016/j.socscimed.2022.115605

Blake, J. (1989) Number of Siblings and Educational Attainment. Science. 245 (4913), 32–36. doi:10.1126/science.2740913

Braudt, D. B., Lawrence, E. M., Tilstra, A. M., Rogers, R. G., & Hummer, R. A. (2019). Family socioeconomic status and early life mortality risk in the United States. Maternal and child health journal, 23, 1382–1391. 10.1007/s10995-019-02799-0

Boerma, J.T. & Bicego, G.T. (1992) Preceding Birth Intervals and Child Survival: Searching for Pathways of Influence. Studies in Family Planning. 23 (4), 243. doi:10.2307/1966886.

Borgerhoff Mulder, M. (1998) Brothers and sisters: How sibling interactions affect optimal parental allocations. Human Nature. 9 (2), 119–161. doi:10.1007/s12110-998-1001-6.

Chapman, S. N., Lahdenperä, M., Pettay, J. E., Lynch, R. F., & Lummaa, V. (2021). Offspring fertility and grandchild survival enhanced by maternal grandmothers in a pre-industrial human society. Scientific Reports, 11(1), 3652. doi:10.1038/s41598-021-83353-3

Chesnais, J.-C. (1992) The demographic transition: stages, patterns, and economic implications : a longitudinal study of sixty-seven countries covering the period 1720-1984. Oxford, Clarendon Press. doi: 10.1093/oso/9780198286592.001.0001

Clutton-brock, T. (2016) Mammal societies. John Wiley & Sons.

Clutton-Brock, T.H. (1988) Reproductive success: studies of individual variation in contrasting breeding systems. University of Chicago Pres.

Corbett, S., Courtiol, A., Lummaa, V., Moorad, J. & Stearns, S. (2018) The transition to modernity and chronic disease: mismatch and natural selection. Nature Reviews Genetics. 19 (7), 419–430. doi:10.1038/s41576-018-0012-3.

Crognier, E., Baali, A. & Hilali, M.-K. (2001) Do ‘helpers at the nest’ increase their parents’ reproductive success? American Journal of Human Biology. 13 (3), 365–373. doi:10.1002/ajhb.1060.

Crognier, E., Villena, M. & Vargas, E. (2002) Helping patterns and reproductive success in Aymara communities. American Journal of Human Biology. 14 (3), 372–379. doi:10.1002/ajhb.10047.

Davatz, J. (1980). Glarner Geschichte für die Schulen des Kantons Glarus. Kantonaler Lehrmittelverlag Glarus.

Drevenstedt, G.L., Crimmins, E.M., Vasunilashorn, S. & Finch, C.E. (2008) The rise and fall of excess male infant mortality. Proceedings of the National Academy of Sciences. 105 (13), 5016–5021. doi:10.1073/pnas.0800221105.

Emlen, S.T. (1995) An evolutionary theory of the family. Proceedings of the National Academy of Sciences. 92 (18), 8092–8099. doi:10.1073/pnas.92.18.8092.

Evans, S.R., Waldvogel, D., Vasiljevic, N. & Postma, E. (2018) Heritable spouse effects increase evolutionary potential of human reproductive timing. Proceedings of the Royal Society B: Biological Sciences. 285 (1876). doi:10.1098/rspb.2017.2763.

Faurie, C., Russell, A.F. & Lummaa, V. (2009) Middleborns Disadvantaged? Testing Birth-Order Effects on Fitness in Pre-Industrial Finns R. Sear (ed.). PLoS ONE. 4 (5), e5680. doi:10.1371/journal.pone.0005680.

Fox, J., Weisberg, S., Price, B., Adler, D. & Bates, D. (2019) car: Companion to Applied Regression. https://CRAN.R-project.org/package=car.

Fox, J., Willführ, K., Gagnon, A., Dillon, L. & Voland, E. (2017) The consequences of sibling formation on survival and reproductive success across different ecological contexts: a comparison of the historical Krummhörn and Quebec populations. History of the Family. 22 (2–3), 364–423. doi:10.1080/1081602X.2016.1193551.

Frayer, D.W. & Wolpoff, M.H. (1985) Sexual Dimorphism. Annual Review of Anthropology. 14, 429–473. doi: https://www.jstor.org/stable/2155603

Fuse, K. & Crenshaw, E.M. (2006) Gender imbalance in infant mortality: A cross-national study of social structure and female infanticide. Social Science & Medicine. 62 (2), 360–374. doi:10.1016/j.socscimed.2005.06.006.

Gillespie, D. O., Russell, A. F., & Lummaa, V. (2013). The effect of maternal age and reproductive history on offspring survival and lifetime reproduction in preindustrial humans. Evolution, 67(7), 1964–1974. doi:10.1111/evo.12078.

Grafen, A. (1988) On the uses of data on lifetime reproductive success. In: T. Clutton-brock (ed.). Reproductive success. University of Chicago Press, Chicago, IL. pp. 454–751.

Hagen, E.H. & Barrett, H.C. (2009) Cooperative Breeding and Adolescent Siblings: Evidence for the Ecological Constraints Model? Current Anthropology. 50 (5), 727–737. doi:10.1086/605328.

Hamilton, W.D. (1964) The genetical evolution of social behaviour. II. Journal of Theoretical Biology. 7 (1), 17–52. doi:10.1016/0022-5193(64)90039-6.

Hartig F (2024). DHARMa: Residual Diagnostics for Hierarchical (Multi-Level / Mixed) Regression Models. R package version 0.4.7, https://CRAN.R-project.org/package=DHARMa

Harvey, P.H. & Clutton-Brock, T.H. (1985) Life History Variation in Primates. Evolution. 39 (3), 559. doi:10.2307/2408653.

Helfrecht, C. & Meehan, C.L. (2016) Sibling effects on nutritional status: Intersections of cooperation and competition across development. American Journal of Human Biology. 28 (2), 159–170. doi:10.1002/ajhb.22763.

Hertwig, R., Davis, J.N. & Sulloway, F.J. (2002) Parental investment: How an equity motive can produce inequality. Psychological Bulletin. 128 (5), 728–745. doi:10.1037/0033-2909.128.5.728.

Hobson, A. (1929) Agricultural Survey of Europe: Switzerland. doi: 10.22004/ag.econ.157084.

Hudson, R. & Trillmich, F. (2008) Sibling competition and cooperation in mammals: challenges, developments and prospects. Behavioral Ecology and Sociobiology. 62 (3), 299–307. doi:10.1007/s00265-007-0417-z.

Invernizzi, L., Lemaître, J. F., & Douhard, M. (2025). The expensive son hypothesis. Journal of Animal Ecology, 94(1), 20–44. doi: 10.1111/1365-2656.14207

Kalmijn, M., & van de Werfhorst, H. G. (2016). Sibship size and gendered resource dilution in different societal contexts. PloS one, 11(8), e0160953.DOI:10.1371/journal.pone.0160953.

Kaplan, H. (1997). The evolution of the human life course. Between Zeus and the salmon: The biodemography of longevity, 175–211.

Kassambara, Alboukadel. (2020) ggpubr: ‘ggplot2’ Based Publication Ready Plots. https://cran.r-project.org/package=ggpubr.

Kramer, K.L. (2005) Children’s Help and the Pace of Reproduction: Cooperative Breeding in Humans. Evolutionary Anthropology: Issues, News, and Reviews. 14 (6), 224–237. doi:10.1002/evan.20082.

Kramer, K.L. (2002) Variation in juvenile dependence: Helping behavior among Maya children. Human Nature. 13 (2), 299–325. doi:10.1007/s12110-002-1011-8.

Kramer, K.L. & Boone, J.L. (2002) Why Intensive Agriculturalists Have Higher Fertility: A Household Energy Budget Approach. Current Anthropology. 43 (3), 511–517. doi:10.1086/340239.

Kubly-Müller, J.J. (1912) Die genealogien-werke des kanotons glarus. In: Archiv für Heraldik. pp. 164–187.

Lack, D. (1947) The significance of clutch-size. Ibis. 89 (2), 302–352. 10.1111/j.1474-919X.1947.tb04155.x.

Lambert, P.S., Zijdeman, R.L., Van Leeuwen, M.H.D., Maas, I. & Prandy, K. (2013) The construction of HISCAM: A stratification scale based on social interactions for historical comparative research. Historical Methods. 46 (2), 77–89. doi:10.1080/01615440.2012.715569.

Lawson, D. W., Alvergne, A., & Gibson, M. A. (2012). The life-history trade-off between fertility and child survival. Proceedings of the Royal Society B: Biological Sciences, 279(1748), 4755–4764.doi:10.1098/rspb.2012.1635

Lee, R. (2003) The Demographic Transition: Three Centuries of Fundamental Change. Journal of Economic Perspectives. 17 (4), 167–190. doi:10.1257/089533003772034943.

Leeuwen, M. van, Maas, I., Miles, A., Edvinsson, S., Karlsson, J., Jarnaes-Erikstad, M.,… & Matthijs, K. (2002). HISCO. Historical international standard classification of occupations (pp. 441-p). Leuven University Press.

Lenth R (2024). emmeans: Estimated Marginal Means, aka Least-Squares Means. R package version 1.10.6, https://CRAN.R-project.org/package=emmeans.

Li, R., Ware, J., Chen, A., Nelson, J. M., Kmet, J. M., Parks, S. E.,… & Perrine, C. G. (2022). Breastfeeding and post-perinatal infant deaths in the United States, a national prospective cohort analysis. The Lancet Regional Health–Americas, 5. doi: 10.1016/j.lana.2021.100094.

Lindström, J. (1999) Early development and fitness in birds and mammals. Trends in Ecology & Evolution. 14 (9), 343–348. doi:10.1016/S0169-5347(99)01639-0.

Lüdecke, D. (2018) ggeffects: Tidy Data Frames of Marginal Effects from Regression Models. Journal of Open Source Software. 3 (26), 772. doi:10.21105/joss.00772.

Lummaa, V. (2001). Reproductive investment in pre–industrial humans: the consequences of offspring number, gender and survival. Proceedings of the Royal Society of London. Series B: Biological Sciences, 268(1480), 1977–1983. doi: 10.1098/rspb.2001.1786.

Marco-Gracia, F.J. & Beltrán Tapia, F.J. (2021) Son Preference, Gender Discrimination, and Missing Girls in Rural Spain, 1750–1950. Population and Development Review. 47 (3), 665–689. doi:10.1111/padr.12406.

Marco-Gracia, F. J., & López-Antón, M. (2025). “For Better, for Worse”: The Role of Siblings in Survival and Biological Well-Being in Rural Aragón (Spain) in the Twentieth Century. Social Science History, 1–29. doi:10.1017/ssh.2025.26

Minge-Kalman, W. (1978) Household Economy during the Peasant-to-Worker Transition in the Swiss Alps. Etnology. 17 (2), 183–196. 10.2307/3773143.

Moucheraud, C., Worku, A., Molla, M., Finlay, J.E., Leaning, J. & Yamin, A.E. (2015) Consequences of maternal mortality on infant and child survival: a 25-year longitudinal analysis in Butajira Ethiopia (1987-2011). Reproductive Health. 12 (S1), S4. doi:10.1186/1742-4755-12-S1-S4.

Muhuri, P.K. & Menken, J. (1997) Adverse Effects of Next Birth, Gender, and Family Composition on Child Survival in Rural Bangladesh. Population Studies. 51 (3), 279–294. doi:10.1080/0032472031000150056.

Nag, M., Peet, R.C., Bardhan, A., Hull, T.H., Johnson, A., Masnick, G.S., Polgar, S., Repetto, R. & Tax, S. (1978) An Anthropological Approach to the Study of the Economic Value of Children in Java and Nepal [and Comments and Reply]. Current Anthropology. 19 (2), 293–306. doi:10.1086/202076.

Nitsch, A., Faurie, C. & Lummaa, V. (2013) Are elder siblings helpers or competitors? Antagonistic fitness effects of sibling interactions in humans. Proceedings of the Royal Society B: Biological Sciences. 280 (1750). doi:10.1098/rspb.2012.2313.

Parker, G.A., Mock, D.W. & Lamey, T.C. (1989) How Selfish Should Stronger Sibs Be? The American Naturalist. 133 (6), 846–868. doi:10.1086/284956.

Pettay, J.E., Helle, S., Jokela, J. & Lummaa, V. (2007) Natural Selection on Female Life-History Traits in Relation to Socio-Economic Class in Pre-Industrial Human Populations. PLoS ONE. 2 (7). doi:10.1371/journal.pone.0000606.

R Core Team (2024). R: A Language and Environment for Statistical Computing. R Foundation for Statistical Computing, Vienna, Austria. https://www.R-project.org/.

Rachoud-Schneider, A.M., Leonhard, M., Schnyder, A., Baumann, W. & Moser, P. (2007) Agriculture. 19 November 2007. Dictionnaire historique de la Suisse (DHS). https://hls-dhs-dss.ch/fr/articles/013933/2007-11-19/ [Accessed: 7 November 2023].

Rickard, I. J. (2008). Offspring are lighter at birth and smaller in adulthood when born after a brother versus a sister in humans. Evolution and Human Behavior, 29, 196–200. 10.1016/j.evolhumbehav.2008.01.006.

Rickard, I.J., Lummaa, V. & Russell, A.F. (2009) Elder brothers affect the life history of younger siblings in preindustrial humans: social consequence or biological cost? Evolution and Human Behavior. 30 (1), 49–57. doi:10.1016/j.evolhumbehav.2008.08.001.

Rickard, I. J., Vullioud, C., Rousset, F., Postma, E., Helle, S., Lummaa, V.,… & Courtiol, A. (2022). Mothers with higher twinning propensity had lower fertility in pre-industrial Europe. nature communications, 13(1), 2886. 10.1038/s41467-022-30366-9

Riswick, T. & Hsieh, Y.-H. (2020) Between rivalry and support: The impact of sibling composition on infant and child mortality in Taiwan, 1906‒1945. Demographic Research. 42, 615–656. doi:10.4054/DemRes.2020.42.21.

Rosenblum, D. (2015) Unintended Consequences of Women’s Inheritance Rights on Female Mortality in India. Economic Development and Cultural Change. 63 (2), 223–248. doi:10.1086/679059.

Roser, M. (2023) Mortality in the past: every second child died. Our World in Data. https://ourworldindata.org/child-mortality-in-the-past.

Schielzeth, H. & Forstmeier, W. (2009) Conclusions beyond support: overconfident estimates in mixed models. Behavioral Ecology. 20 (2), 416–420. doi:10.1093/beheco/arn145.

Sear, R. (2008) Kin and Child Survival in Rural Malawi: Are Matrilineal Kin Always Beneficial in a Matrilineal Society? Human Nature. 19 (3), 277–293. doi:10.1007/s12110-008-9042-4.

Sear, R. & Mace, R. (2008) Who keeps children alive? A review of the effects of kin on child survival. Evolution and Human Behavior. 29 (1), 1–18. doi:10.1016/j.evolhumbehav.2007.10.001.

Sear, R., Steele, F., Mcgregor, I.A. & Mace, R. (2002) The effects of kin on child mortality in rural gambia. Demography. 39 (1), 43–63. doi: 10.1353/dem.2002.0010

Ritzmann-Blickenstorfer, H. (Ed.). (1996). Historische Statistik der Schweiz: Statistique historique de la Suisse. Chronos.

Singer, J.D. & Willett, J.B. (2003). Applied Longitudinal Data Analysis: Modeling Change and Event Occurrence. 1st edition. Oxford University Press New York. doi:10.1093/acprof:oso/9780195152968.001.0001.

Steelman, L.C., Powell, B., Werum, R. & Carter, S. (2002) Reconsidering the Effects of Sibling Configuration: Recent Advances and Challenges. Annual Review of Sociology. 28 (1), 243–269. doi:10.1146/annurev.soc.28.111301.093304.

Suanet, B. & Bras, H. (2014) Sibling Position and Marriage Timing in the Netherlands, 1840–1922: A Comparison across Social Classes, Local Contexts, and Time. Journal of Family History. 39 (2), 126–139. doi:10.1177/0363199013506986.

Thurstans, S., Opondo, C., Seal, A., Wells, J.C., Khara, T., Dolan, C., Briend, A., Myatt, M., Garenne, M., Mertens, A., Sear, R. & Kerac, M. (2022) Understanding Sex Differences in Childhood Undernutrition: A Narrative Review. Nutrients. 14 (5), 948. doi:10.3390/nu14050948.

Trillmich, F. & Wolf, J.B.W. (2008) Parent–offspring and sibling conflict in Galápagos fur seals and sea lions. Behavioral Ecology and Sociobiology. 62 (3), 363–375. doi:10.1007/s00265-007-0423-1.

Trivers, R.L. (1974) Parent-Offspring Conflict. American Zoologist. 14 (1), 249–264. 10.1093/icb/14.1.249

Voland, E. & Dunbar, R.I.M. (1995) Resource competition and reproduction: The relationship between economic and parental strategies in the Krummhörn population (1720–1874). Human Nature. 6 (1), 33–49. doi:10.1007/BF02734134.

Volk, A.A. & Atkinson, J.A. (2013) Infant and child death in the human environment of evolutionary adaptation. Evolution and Human Behavior. 34 (3), 182–192. doi:10.1016/j.evolhumbehav.2012.11.007.

Whitworth, A. & Stephenson, R. (2002) Birth spacing, sibling rivalry and child mortality in India. Social Science & Medicine. 55 (12), 2107–2119. doi:10.1016/S0277-9536(02)00002-3.

Wickham, H. (2016) ggplot2: Elegant Graphics for Data Analysis. https://ggplot2.tidyverse.org.

World Health Organization. (2022, January, 28). Child mortality (under 5 years). WHO. https://www.who.int/news-room/fact-sheets/detail/child-mortality-under-5-years

Zhou, L., Ge, E., Micheletti, A.J.C., Chen, Y., Du, J. & Mace, R. (2022) Monks relax sibling competition over parental resources in Tibetan populations A. Ridley (ed.). Behavioral Ecology. 33 (6), 1070–1079. doi:10.1093/beheco/arac059.

