## Supplementary material for "Age- and sex-dependent sibling effects on early-life survival in preindustrial humans": All supplementary tables

**Table S1.** Logistic GLMM showing the association between childhood survival and the number of older siblings from *m1* (N = 2941). Here, we look at the total number of older siblings of the focal individual and present their associations with childhood survival upon the other factors. Odds ratios, 95% confidence intervals, variation explained by random effects and p-values from LRT tests are presented. Interactions are shown with a multiplication symbol (×).

| | **Childhood survival (<5)** | | |
|---|---|---|---|
| Fixed effects | Odds ratio | 95% CI | P-value |
| *Intercept* | **3.645 **** | **1.37-9.67** | |
| Sex (male) | 0.868 | 0.729-1.033 | 0.113 |
| The number of older siblings | 0.910 | 0.767-1.079 | 0.282 |
| Mother age | 0.908 | 0.755-1.091 | 0.304 |
| Mother age$^2$ | **0.891 **** | **0.823- 0.965** | **0.005** |
| Father age | **1.206 *** | **1.022- 1.422** | **0.027** |
| Maternal survival | **1.923 *** | **1.068- 3.461** | **0.032** |
| Paternal survival | 0.650 | 0.299- 1.415 | 0.268 |
| Socioeconomic status | 0.934 | 0.840- 1.038 | 0.209 |
| Parish (linthal) | **0.674 **** | **0.527- 0.861** | **0.003** |
| The number of grandparents living at birth | 0.979 | 0.871- 1.099 | 0.717 |
| Random effects | Variance | P-value | |
| Cohort (intercept) | 0.193 | 0.120 | |
| Family (intercept) | **0.559 ***** | **<0.001** | |
| The number of older siblings\| Family (random slope) | 0.258 | 0.157 | |

**Table S2.** Logistic GLMM showing the association between childhood survival and the number of older siblings from *m1* including all relevant interactions (N = 2941). Here, we look at the total number of older siblings of the focal individual and present their associations with childhood survival upon the other factors. Odds ratios, 95% confidence intervals, variation explained by random effects and p-values from LRT tests are presented. Interactions are shown with a multiplication symbol (×).

| | **Childhood survival (<5)** | | |
|---|---|---|---|
| Fixed effects | Odds ratio | 95% CI | P-value |
| *Intercept* | 3.950 | 1.352-9.535 | |
| Sex (male) | 0.867 | 0.728-1.032 | |
| The number of older siblings | 0.885 | 0.730-1.073 | |
| Mother age | 0.909 | 0.757-1.093 | |
| Mother age² | 0.892 | 0.824- 0.965 | |
| Father age | 1.206 | 1.022- 1.423 | |
| Maternal survival | 1.931 | 1.073- 3.475 | |
| Paternal survival | 0.656 | 0.301- 1.430 | |
| Socioeconomic status | 0.934 | 0.840- 1.038 | |
| Parish (linthal) | 0.675 | 0.528- 0.863 | |
| The number of grandparents living at birth | 0.980 | 0.872- 1.100 | |
| Sex (male) × the number of older siblings | 1.054 | 0.887 - 1.254 | 0.551 |
| Random effects | Variance | P-value | |
| Cohort (intercept) | 0.193 | | |
| Family (intercept) | 0.559 | | |
| The number of older siblings\| Family (random slope) | 0.257 | | |

**Table S3.** Logistic GLMM showing the association between childhood survival and the number of older siblings from *m2* (N = 2941). Here, we have decomposed the number of older siblings into deceased and living older siblings of the focal individual and present their associations with childhood survival upon the other factors. Odds ratios, 95% confidence intervals, variation explained by random effects and p-values from LRT tests are presented. Interactions are shown with a multiplication symbol (×).

| | **Childhood survival (<5)** | | |
|---|---|---|---|
| Fixed effects | Odds ratio | 95% CI | P-value |
| *Intercept* | **3.653 **** | **1.397-9.552** | |
| Sex (male) | 0.886 | 0.745-1.055 | |
| The number of living older siblings | 0.947 | 0.792-1.131 | |
| The number of deceased older siblings | 1.121 | 0.947-1.325 | |
| Mother age | **0.831 *** | **0.697-0.991** | **0.039** |
| Mother age$^2$ | **0.888 **** | **0.821-0.959** | **0.003** |
| Father age | 1.151 • | 0.979-1.353 | 0.088 |
| Maternal survival | **1.901 *** | **1.067-3.387** | **0.031** |
| Paternal survival | 0.656 | 0.304-1.415 | 0.272 |
| Socioeconomic status | 0.929 | 0.837-1.032 | 0.172 |
| Parish (linthal) | **0.664 **** | **0.519-0.849** | **0.002** |
| The number of grandparents living at birth | 0.982 | 0.876-1.102 | 0.763 |
| Sex (male) × the number of living older siblings | **1.252 *** | **1.040-1.508** | **0.018** |
| Sex (male) × the number of deceased older siblings | **0.817 *** | **0.684- 0.975** | **0.026** |
| Random effects | Variance | P-value | |
| Cohort (intercept) | 0.193 | 0.110 | |
| Family (intercept) | **0.560** | **<0.001***** | |
| The number of living older siblings\| Family (random slope) | 0.000 | 1.000 | |
| The number of deceased older siblings\| Family (random slope) | 0.000 | 1.000 | |

**Table S4.** Logistic GLMM showing the association between childhood survival and the number of older siblings from *m3* (N = 2941). Here, we have decomposed the number of older siblings into deceased and living older siblings born close and far in age (≥5) of the focal individual and present their associations with childhood survival upon the other factors. Odds ratios, 95% confidence intervals, variation explained by random effects and p-values from LRT tests are presented. Interactions are shown with a multiplication symbol (×).

| | **Childhood survival (<5)** | | |
|---|---|---|---|
| Fixed effects | Odds ratio | 95% CI | P-value |
| *Intercept* | **3.607 **** | **1.376-9.453** | |
| Sex (male) | 0.878 | 0.738-1.045 | |
| The number of living older ≥5 siblings | 1.056 | 0.900-1.238 | 0.516 |
| The number of living older <5 siblings | 0.883 | 0.774-1.007 | |
| The number of deceased older siblings | 1.017 | 0.881-1.174 | 0.817 |
| Mother age | **0.830 *** | **0.694-0.993** | **0.041** |
| Mother age² | **0.881 **** | **0.811-0.957** | **0.003** |
| Father age | 1.148 • | 0.975-1.353 | 0.098 |
| Maternal survival | **1.949 *** | **1.091-3.482** | **0.026** |
| Paternal survival | 0.658 | 0.305-1.421 | 0.278 |
| Socioeconomic status | 0.929 | 0.836-1.033 | 0.176 |
| Parish (linthal) | **0.665 **** | **0.518-0.853** | **0.002** |
| The number of grandparents living at birth | 0.9855 | 0.878-1.106 | 0.806 |
| Sex (male) × the number of living older <5 siblings | **1.314 **** | **1.100-1.569** | **0.003** |
| Random effects | Variance | P-value | |
| Cohort (intercept) | 0.198 • | 0.096 | |
| Family (intercept) | **0.575 ***** | **<0.001** | |
| The number of living older ≥5 siblings\| Family (random slope) | 0 | 1 | |
| The number of living older <5 siblings\| Family (random slope) | 0 | 1 | |

| | | |
|---|---|---|
| The number of deceased older siblings\| Family (random slope) | 0 | 1 |

**Table S5.** Logistic GLMM showing the association between childhood survival and the number of older siblings from *m3* including all relevant (N = 2941). Here, we have decomposed the number of older siblings into deceased and living older siblings born close and far in age (≥5) of the focal individual and present their associations with childhood survival upon the other factors. Odds ratios, 95% confidence intervals, variation explained by random effects and p-values from LRT tests are presented. Interactions are shown with a multiplication symbol (×).

| | **Childhood survival (<5)** | | |
|---|---|---|---|
| Fixed effects | Odds ratio | 95% CI | P-value |
| *Intercept* | 3.652 | 1.392-9.585 | |
| Sex (male) | 0.888 | 0.746-1.057 | |
| The number of living older ≥5 siblings | 1.016 | 0.848-1.218 | |
| The number of living older <5 siblings | 0.889 | 0.779-1.014 | |
| The number of deceased older siblings | 1.099 | 0.929-1.301 | |
| Mother age | 0.829 | 0.693-0.990 | |
| Mother age$^2$ | 0.881 | 0.811-0.957 | |
| Father age | 1.146 | 0.973-1.349 | |
| Maternal survival | 1.922 | 1.076-3.433 | |
| Paternal survival | 0.654 | 0.303-1.414 | |
| Socioeconomic status | 0.928 | 0.835-1.031 | |
| Parish (linthal) | 0.666 | 0.520-0.851 | |
| The number of grandparents living at birth | 0.984 | 0.877-1.104 | |
| Sex (male) × the number of living older <5 siblings | **1.302** ** | **1.089-1.557** | **0.004** |
| Sex (male) × the number of living older ≥5 siblings | 1.088 | 0.900-1.315 | 0.386 |
| Sex (male) × the number of deceased older siblings | 0.850 • | 0.709-1.019 | 0.080 |

| Random effects | Variance | P-value |
| --- | --- | --- |
| Cohort (intercept) | 0.191 | |
| Family (intercept) | 0.566 | |
| The number of living older ≥5 siblings\| Family (random slope) | 0 | |
| The number of living older <5 siblings\| Family (random slope) | 0 | |
| The number of deceased older siblings\| Family (random slope) | 0 | |

**Table S6.** Logistic GLMM showing the association between childhood survival and the number of older siblings from *m4* including all relevant interactions (N = 2941). Here, we have decomposed the number of older siblings into deceased and living older sisters and brothers born close and far in age (≥5) of the focal individual and present their associations with childhood survival upon the other factors. Odds ratios, 95% confidence intervals, variation explained by random effects and p-values from LRT tests are presented. Interactions are shown with a multiplication symbol (×).

| | **Childhood survival (<5)** | | |
|---|---|---|---|
| Fixed effects | Odds ratio | 95% CI | P-value |
| *Intercept* | 3.730 | 1.383-10.061 | |
| Sex (male) | 0.870 | 0.729-1.040 | |
| The number of living older ≥5 brothers | 1.050 | 0.888-1.242 | |
| The number of living older <5 brothers | 0.798 | 0.699-0.912 | |
| The number of deceased older brothers | 1.072 | 0.918-1.251 | |
| The number of living older ≥5 sisters | 1.001 | 0.843-1.187 | |
| The number of living older <5 sisters | 1.039 | 0.910-1.186 | |
| The number of deceased older sisters | 1.080 | 0.925-1.261 | |
| Mother age | 0.824 | 0.686-0.989 | |
| Mother age² | 0.870 | 0.798-0.948 | |
| Father age | 1.146 | 0.969-1.355 | |
| Maternal survival | 2.071 | 1.136-3.775 | |
| Paternal survival | 0.630 | 0.286-1.389 | |
| Socioeconomic status | 0.924 | 0.829-1.030 | |
| Parish (linthal) | 0.655 | 0.508-0.843 | |
| The number of grandparents living at birth | 0.985 | 0.876-1.108 | |
| Sex (male) × the number of living older ≥5 brothers | 1.167 | 0.952- 1.432 | 0.140 |
| Sex (male) × the number of living older <5 brothers | **1.288 **** | **1.077- 1.539** | **0.006** |
| Sex (male) × the number of deceased older brothers | 0.888 | 0.742- 1.061 | 0.192 |

| | | | |
|---|---|---|---|
| Sex (male) × the number of living older ≥5 sisters | 0.927 | 0.755- 1.139 | 0.475 |
| Sex (male) × the number of living older <5 sisters | 1.160 | 0.964- 1.396 | 0.124 |
| Sex (male) × the number of deceased older sisters | 0.920 | 0.765- 1.107 | 0.381 |
| Random effects | Variance | P-value | |
| Cohort (intercept) | 0.200 | | |
| Family (intercept) | 0.581 | | |
| The number of living older ≥5 brothers\| Family (random slope) | 0 | | |
| The number of living older <5 brothers\| Family (random slope) | 0 | | |
| The number of deceased older brothers\| Family (random slope) | 0 | | |
| The number of living older ≥5 sisters\| Family (random slope) | 0.268 | | |
| The number of living older <5 sisters\| Family (random slope) | 0 | | |
| The number of deceased older sisters\| Family (random slope) | 0 | | |
